# Identification of natural Zika virus peptides presented on the surface of paediatric brain tumour cells by HLA class I

**DOI:** 10.1101/2024.09.17.613406

**Authors:** Matt Sherwood, Ben Nicholas, Alistair Bailey, Thiago G. Mitsugi, Carolini Kaid, Oswaldo K. Okamoto, Paul Skipp, Rob M. Ewing

**Author notes:** **Correspondence to:** Professor Rob Ewing, B85, Life Sciences Building, University of Southampton, University Road, Highfield, Southampton, Hants., SO17 1BJ. Equal contributions (Oswaldo K. Okamoto, Paul Skipp, Rob M. Ewing).

## Abstract

Despite decades of research, survival from brain cancer has scarcely improved and is drastically lower than that of other cancers. Novel therapies, such as immunotherapy, hold great promise for treating brain tumours and are desperately needed. Zika virus (ZIKV) infects and kills aggressive cancer cells with stem-like properties (CSCs) from both paediatric and adult brain tumours. Whilst T cell recruitment into ZIKV-infected brain tumours is becoming well documented, the specific mechanisms through which they are activated are poorly understood. We address this by employing a combined LC-MS/MS global proteome and immunopeptidome approach to describe, for the first time, human leukocyte antigen (HLA) presentation of ZIKV peptides on the surface of infected brain tumour cells. We first show that HLA class I (HLA-I) antigen processing & presentation is the most highly enriched immune response pathway in the global proteome of aggressive paediatric USP7-ATRT brain tumour cells following ZIKV infection. We identify USP7-ATRT cells as a good immunopeptidome model due to their homozygous of the globally most common HLA-A allotype (A*02:01). We predict the majority of the 19 ZIKV peptides that we identify here to strongly bind and be presented by HLA-A*02:01. We show that immunopeptide presentation corresponds with cellular ZIKV protein abundance, with ten peptides arising from the most abundant viral protein; non-structural protein 3 (NS3). Specifically, we show the ZIKV NS3 helicase domain to be a rich source of peptides. Finally, we verify that the identified ZIKV peptides do not mimic predicted peptides of the human proteome. The ZIKV peptides we identify here are potential candidates for developing novel epitope-specific brain tumour immunotherapies, and our findings provide potential insight into the efficacious cytotoxic T cell response that oncolytic ZIKV virotherapy can induce against brain tumours.

**Author Summary:** Viruses can attack cancer through two mechanisms: 1) infecting and killing the cancer cell and 2) activating the immune system against the tumour. Zika virus (ZIKV) uses both mechanisms to fight brain cancer. Here, we employ a powerful proteomic technique to identify fragments of viral proteins (peptides) presented by cell surface receptors on brain cancer cells infected with ZIKV. In the human body, immune system cells such as T cells recognise and become activated in response to these viral peptides and subsequently attack the infected patient tumour. We identify 19 ZIKV peptides, three of which have been shown previously to elicit T cell responses, four identified elsewhere, and twelve are novel. Our work helps delineate a component of how ZIKV acts as an immunotherapy, the T cell-specific immune response that the virus raises to promote clearance of brain tumours. The significance of our study is that the ZIKV peptides we identify may lead to the development of a novel brain tumour immunotherapy.

## Introduction

Central nervous system (CNS) tumours account for approximately one-fifth of all childhood cancer cases and disproportionately are the largest cause of cancer-related mortality in children (1,2). These tumours exhibit high lethality, and the aggressive nature of standard-of-care therapy often leaves survivors with severe sequelae that significantly affect their quality of life. CNS tumours are commonly immunosuppressed, and there is significant interest in developing immunotherapeutic strategies to circumvent this suppression by activating a patient’s immune system against their tumour (3).

Oncolytic virotherapy, a specific class of immunotherapy, exploits oncolytic viruses (OVs) that preferentially infect and destroy tumour cells, with minimal pathology against non-cancerous cells and tissue. These viruses present a unique advantage in targeting highly heterogeneous and immunosuppressive cancers, such as CNS tumours, as the second pillar of oncolytic virotherapy is the ability to mount an anti-tumoral immune response. Oncolytic virotherapy clinical studies have generally reported low toxicity and minimal adverse effects in patients, and there are >200 clinical trials underway to treat aggressive forms of cancer using OVs (4–6). Recently, the oncolytic herpes virus G47Δ was approved in Japan for glioblastoma treatment; the first oncolytic virotherapy against any nervous system tumour in the clinic (7). CNS tumours may generate a suppressive tumour immune microenvironment (TIME) through a combination of intrinsic (reduced HLA antigen presentation, immune checkpoint blockade and immunosuppressive cytokine secretion) and extrinsic (immunosuppressive immune cell recruitment) factors (8). OVs can remodel these factors to overcome the immunosuppressive TIME (9). Due to this remodelling, there is significant interest in employing OVs as adjuvants to other immunotherapies, including monoclonal antibodies, CAR-T cells, cancer vaccines, checkpoint inhibitors and small molecule inhibitors (10).

ZIKV is neuropathogenic and causes congenital ZIKV syndrome (CZS) in 5-14% of babies born to women who contract ZIKV during pregnancy and pass the virus to the fetus via vertical transmission (11). ZIKV infects and diminishes the pool of fetal neural stem and progenitor cells (NPCs) through induction of differentiation or cell death, subsequently leading to the underdevelopment of the fetal brain (12–15). In contrast, ZIKV infection in children is mild, and only 1:5 people are symptomatic. ZIKV infection is generally self-limiting as symptoms resolve within a week or less, and the majority of symptomatic children present only with flu-like symptoms (11,16). This mild infection in children, in cohort with ZIKV’s neurotropism, highlights the virus as a promising candidate to employ against paediatric CNS tumours.

Since 2017, members of our research team and others have demonstrated that ZIKV infects and induces oncolysis of paediatric brain tumour cells *in vitro* and *in vivo*, and mounts an immune response against spontaneous brain tumours in canines (17–19). ZIKV stimulates the infiltration of multiple immune cell types into CNS tumours, including CD8+ and CD4+ T cells, which contribute to ZIKV-induced tumour clearance (20–22). This branch of the adaptive immune response is brought about by viral peptide presentation by the HLA class I and II protein complexes on the cell surface to CD8+ and CD4+ T cells, respectively. Whilst the recruitment of these immune cells into ZIKV-infected CNS tumours is becoming well documented, their cognate HLA-presented ZIKV peptides remain unknown.

Previously, we demonstrated the aggressive paediatric atypical teratoid rhabdoid tumour (ATRT) cell line, USP7-ATRT, to have CSC properties and to be highly susceptible to ZIKV infection and oncolysis (18). In the present study, we show ZIKV infection to enrich MHC Class I antigen processing and presentation at the proteome level in these paediatric brain tumour cells. To investigate this response further, we perform HLA typing to show that USP7 cells are homozygous for all three classical HLA alleles and express the globally most common HLA-A allotype (A*02:01). Performing immunopeptidome profiling, we identify a specific list of 19 ZIKV peptides from infected USP7 cells, predicted to be processed and presented by HLA-I molecules. The mass spectrometry proteomics and analysis used here are similar to our recent influenza work (23). To our knowledge, here we document ZIKV epitopes presented by human CNS tumour cells for the first time. We provide new ZIKV epitopes as novel targets for immunotherapy, and their identification should lead to future work that facilitates understanding of how the immune response can be coupled with the ZIKV oncolytic response.

## Results

### ZIKV infection enriches HLA Class I presentation pathways

We first investigated if ZIKV infection moderates innate or adaptive immune responses within the aggressive paediatric brain tumour cell line USP7. Global proteome analysis and plotting the top and bottom 50 ranked proteins of ZIKV-infected USP7 cells identify the ZIKV polyprotein as one of the top-ranked proteins at 24 hours post-infection (hpi) (Figure 1A). Considering the ranked protein list for immune response proteins, 20% of the top 30 proteins (HUWE1, LTN1, PSMB5, PSMD1, RNF213 and SEC61G) are involved in antigen processing and presentation by MHC Class I (Figure 1A). Performing Gene Set Enrichment Analysis (GSEA) with an innate and adaptive immune response Reactome database, we identify ZIKV infection to significantly enrich both innate and adaptive responses, with “Class I MHC mediated antigen processing & presentation” of the adaptive immune response being the only significantly enriched subpathway in the infected samples (Figure 1B and 1C). To determine if this response is also observed for adult brain tumours, we repeated our GSEA analysis on a pre-clinical RNA-Seq dataset from eight glioblastoma (GBM) patients comparing ZIKV-mCherry positive versus negative primary cell populations (Figure 1D). As for ZIKV-infected USP7, “Class I MHC mediated antigen processing & presentation” was the most significantly enriched immune response term in the ZIKV positive adult brain tumour cells (Figure 1D). To conclude, ZIKV infection enriches antigen processing and presentation by MHC Class I in paediatric and adult brain tumour cells at the proteome and transcriptome levels, respectively. We sought to investigate this further.

**Figure 1:**
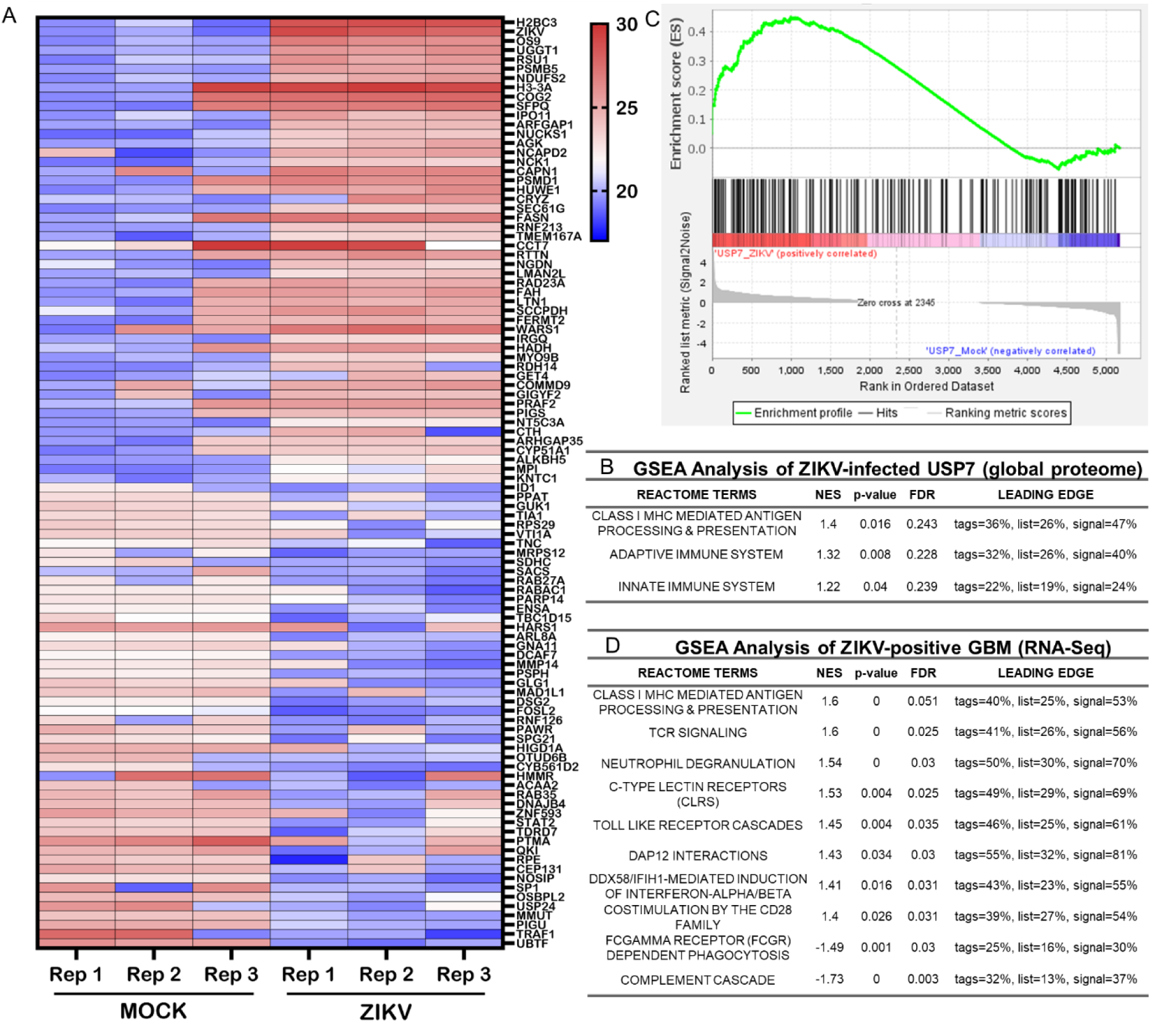
ZIKV infection enriches the HLA Class I pathway in brain tumour cells. **(A)** Log2(LFQ intensity) heatmap of top and bottom 50 ranked proteins from mock and ZIKV-infected USP7 global proteome. GSEA enrichment for Reactome innate and adaptive immune system pathways in **(B)** ZIKV-infected USP7 cells (n=3) and **(D)** ZIKV-positive primary glioblastoma cells (n=8). **(C)** GSEA Class I MHC mediated antigen processing & presentation enrichment plot for ZIKV-infected USP7 cells. Normalised enrichment score (NES) denotes the degree to which the enrichment increased (+) or decreased (−) (p ≤ 0.05, FDR ≤ 0.25).

### USP7 cells predominately present HLA-A*02:01 peptides

Prior to investigating whether infection led to ZIKV-derived HLA immunopeptide presentation by USP7 cells, we first sought to understand what HLA allotypes USP7 cells express and what peptides these HLA molecules present. HLA typing of bulk RNA-Seq of USP7 cells identifies that they are homozygous for all three classical HLA alleles (Table 1). USP7 cells express the globally most common HLA-A allotype (HLA-A*02:01), thus presenting them as a good immunopeptidome model due to their high population coverage (Table 1) (24). Performing immunopeptidomics, we assessed the class I and II HLA immunopeptidomes of mock and ZIKV-infected USP7 cells (Table 2). Supporting our observations at the global proteome level, we predominantly observe peptides with lengths consistent with the nine amino acid (aa) preference for presentation by HLA-I rather than the longer peptides presented by HLA-II (Figure 2A). As expected for non-professional antigen-presenting cells, no significant numbers of HLA-II immunopeptides were recovered (Table 2). ZIKV infection had no effect on the length distribution of the peptides presented (Figure 2A). Unbiased cluster analysis of all the distinct observed 9-mer peptides from ZIKV-infected USP7 cells identified 74%, 14%, and 12% of the 9-mer peptides to be presented by HLA-A*02:01, HLA-B*44:02 and HLA-C*05:01, respectively (Figure 2B). These are consistent with the USP7 HLA-I allotypes identified by HLA typing (Table 1). Peptide length distributions from ZIKV-infected USP7 cells show a dominance of 9-mer peptides, with differing minor populations of 8, 10 or 11-mer peptides, across the three classical HLA molecules (Figure 2C). To conclude, USP7 cells predominantly present 9-mer peptides by the three classical HLA molecules, with nearly three-quarters presented by the globally most common HLA-A allotype HLA-A*02:01.

**Figure 2:**
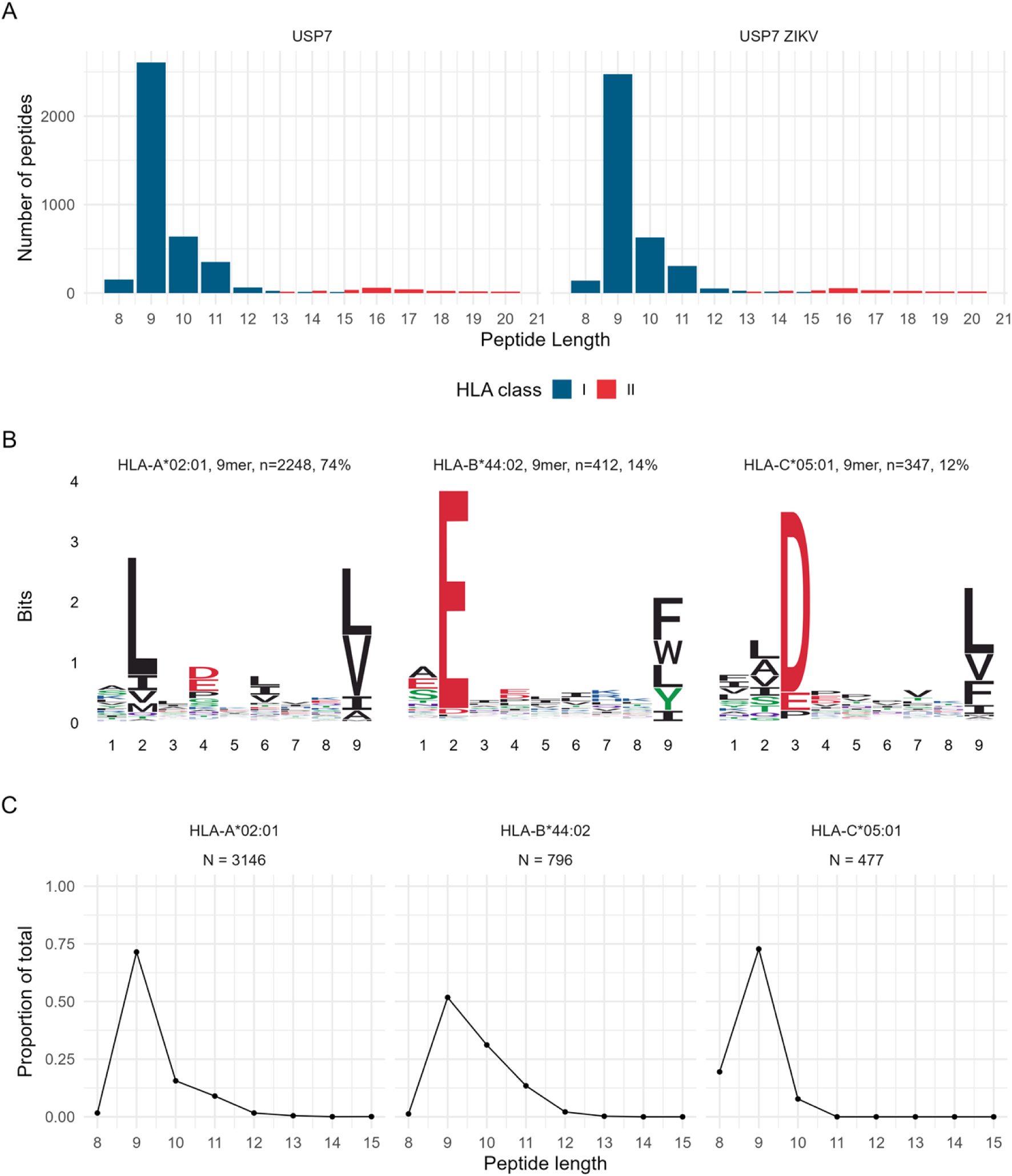
The immunopeptidomes of USP7 brain tumour cells. **(A)** Length distribution of HLA-I and II immunopeptides presented by mock or ZIKV-infected USP7 cells. HLA-I peptides in blue and HLA-II peptides in red. **(B)** Class I HLA allotype 9-mer binding motifs derived from ZIKV-infected USP7 immunopeptides by unsupervised clustering using MixMHCp. **(C)** Length distributions of peptides from ZIKV-infected USP7 cells according to clustered HLA-I allotype.

**Table 1:** USP7 cell HLA allotypes.

| Locus | HLA allotype<sup>1</sup> |
| --- | --- |
| HLA-A | A*02:01 |
| HLA-B | B*44:02 |
| HLA-C | C*05:01 |
| HLA-DRB1 | DRB1*15:01 |

**Table 2:**
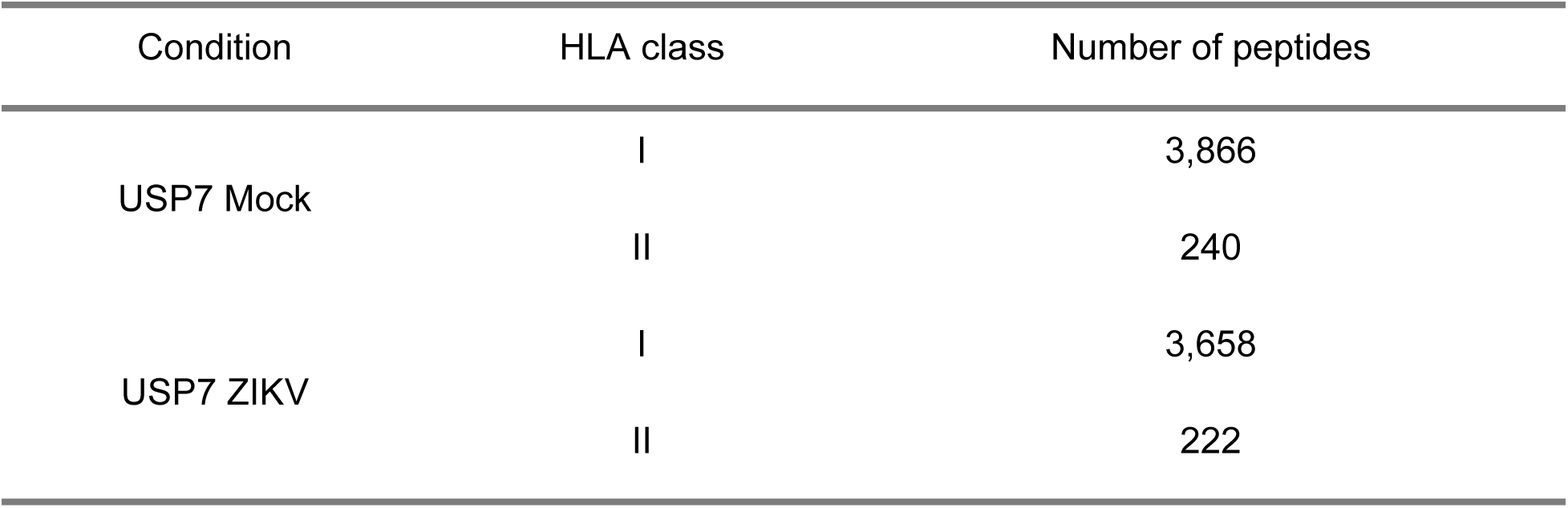
Number of USP7 immunopeptides.

| Condition | HLA class | Number of peptides |
| --- | --- | --- |
| USP7 Mock | I | 3,866 |
|  | II | 240 |
| USP7 ZIKV | I | 3,658 |
|  | II | 222 |

### USP7 cells present ZIKV HLA-I immunopeptides, and presentation corresponds with protein abundance in the HLA-I pathway

We next investigated whether infection led to ZIKV-derived HLA immunopeptide presentation and identify 19 HLA-I peptides derived from six of the ten ZIKV proteins (Table 3). Consistent with the low HLA-II expression by non-professional antigen-presenting cells, no HLA-II presented ZIKV peptides were observed. To examine which HLA-I allotypes the observed ZIKV peptides were likely presented by, and to place these observations in the context of the host cell 9,10 and 11-mer peptides, we predicted ZIKV peptide binding affinities using NetMHC (Figure 3A, Table 4). HLA-A*02:01, HLA-B*44:02 and HLA-C*05:01 were predicted to bind and present twelve, six and one ZIKV peptides, respectively (Table 4). For HLA-A*02:01, all peptides are strong binders, and at least one of the 9, 10 and 11-mers are within the top three predicted binding peptides (Figure 3A). For HLA-B*44:02, the ZIKV NS5 9-mer is the top predicted binding peptide, all four ZIKV 10-mers are within the top seven predicted binding peptides, and the ZIKV NS3 11-mer is a low-affinity binder (Figure 3A). For HLA-C*05:01, the ZIKV NS3 9-mer is a low-affinity binder and was not plotted (Table 4). This data indicates that 17 of the identified ZIKV peptides are high-affinity binding peptides and are commonly within the top predicted binding peptides for each HLA allotype. Interestingly, we observe most ZIKV immunopeptides to be derived from the ZIKV NS3 followed by the NS5 RNA-dependent RNA polymerase (Table 3). To investigate the potential reason for this, we plotted the protein abundances of the ten ZIKV proteins and observed NS3 as the most abundant ZIKV protein, followed by NS5 (Figure 3B). When protein abundances are considered alongside the ZIKV peptide number, a clear relationship emerges between the two (Figure 3B, Table 3). ZIKV NS3 and NS5 are the most abundant proteins and yield the most immunopeptides, therefore indicating that HLA-I presentation corresponds with protein abundance within its peptide processing pathway. This is further supported by the absence of immunopeptides derived from ZIKV Envelope and Membrane proteins, as HLA-I peptides are predominately derived from intracellular cytosolic proteins (25). To conclude, HLA-A*02:01 and HLA-B*44:02 specific high-affinity ZIKV peptide presentation occurs on the surface of USP7 cells, and peptide presentation corresponds with protein abundance in the HLA-I pathway.

**Figure 3:**
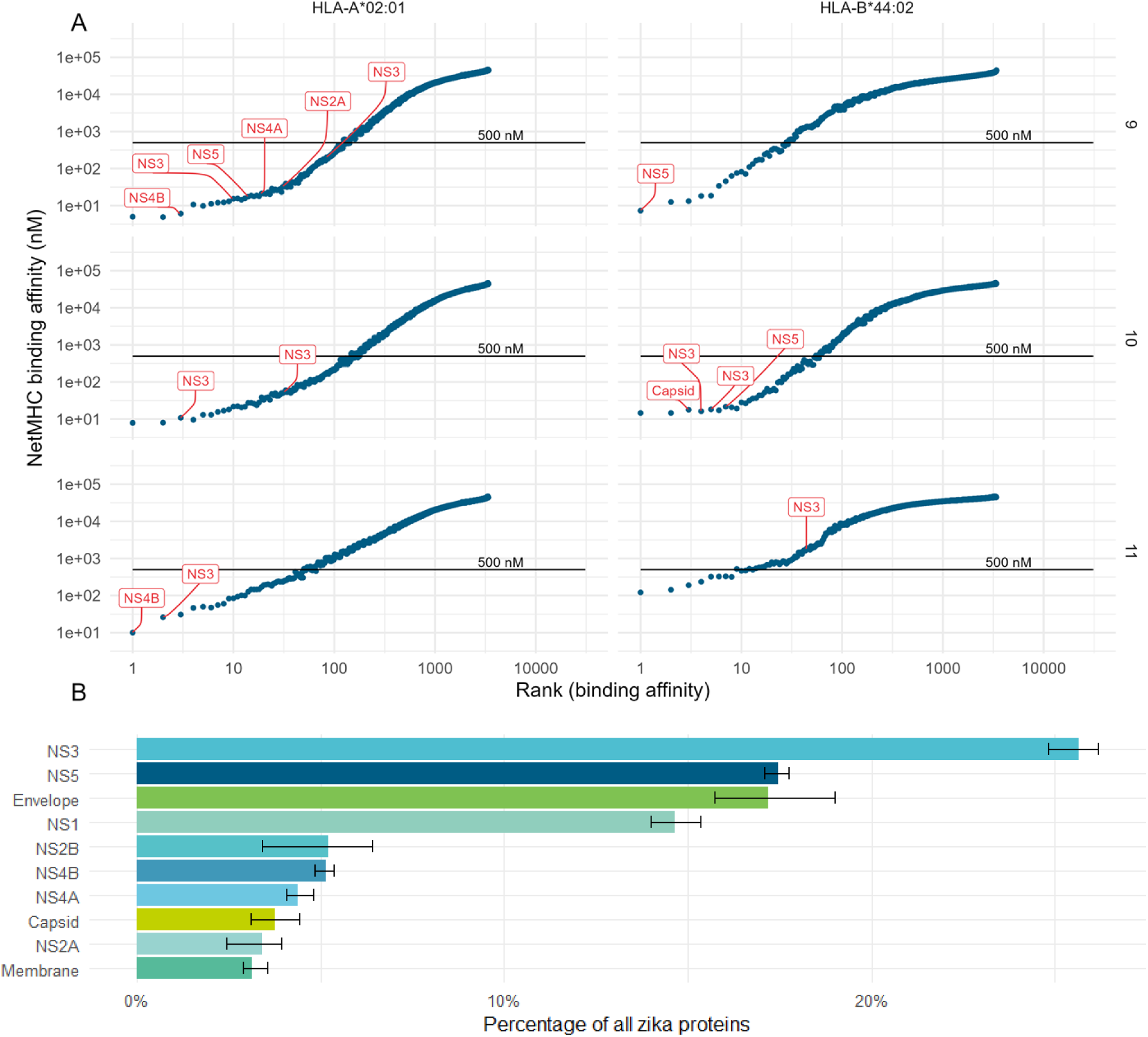
ZIKV immunopeptides binding affinity and viral protein abundance. **(A)** NetMHC binding predictions of all possible 9,10 and 11-mer ZIKV peptides (rows) to the USP7 HLA-I allotypes (columns). HLA binding affinity (y-axis) is plotted against the peptides rank (x-axis). Observed peptides and their source protein are indicated in red, with the black line indicating the 500 nM threshold below which a peptide is considered a strong binder. **(B)** Barplot of the proportion of ZIKV proteins observed in the global proteome of ZIKV-infected USP7 cells.

**Table 3:** Number of USP7 ZIKV immunopeptides per protein.

| Protein | Number of immunopeptides |
| --- | --- |
| NS3 | 10 |
| NS5 | 4 |
| NS4B | 2 |
| Capsid | 1 |
| NS2A | 1 |
| NS4A | 1 |

**Table 4:**
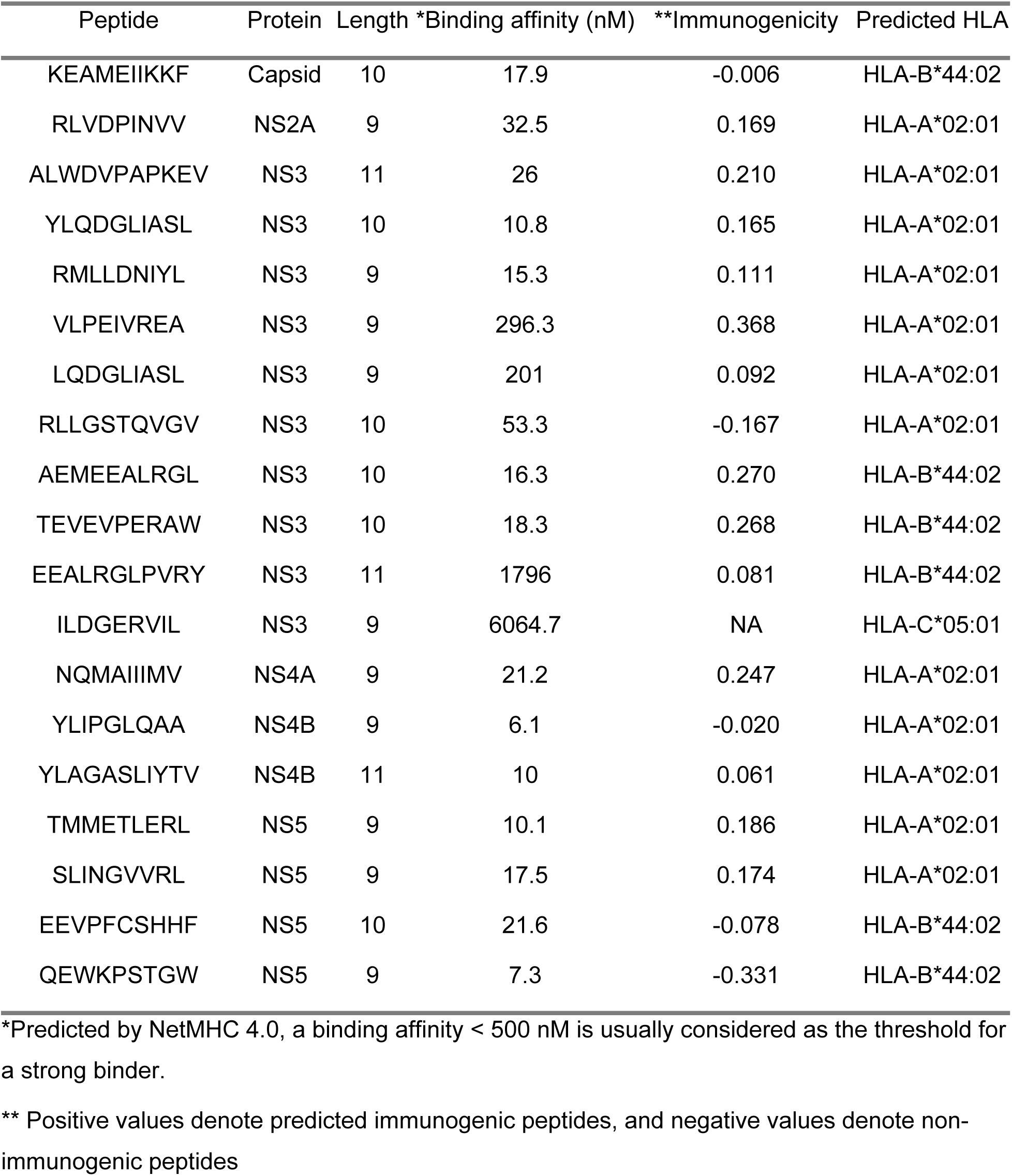
USP7 ZIKV immunopeptides.

### The ZIKV NS3 helicase is a rich source of immunopeptides

Mapping the 19 ZIKV peptides onto the ZIKV polyprotein identifies two peptide-rich regions in the NS3 helicase domain and one in NS5 (Figure 4). For NS3, RMLLDNIYL, YLQDGLIASL, and LQDGLIASL overlap and reside within the continuous 17aa sequence RMLLDNIYLQDGLIASL, whilst AEMEEALRGL and EEALRGLPVRY overlap and reside within the continuous 14aa sequence AEMEEALRGLPVRY. For NS5, QEWKPSTGW and EEVPFCSHHF are non-overlapping and reside within the continual 22aa sequence QEWKPSTGWDNWEEVPFCSHHF. Making *in silico* immunogenicity predictions of the 18 identified HLA-A*02:01 and HLA-B*44:02 presented ZIKV peptides identifies 13 as immunogenic and five as non-immunogenic (Table 4). Interestingly, all eight ZIKV NS3 helicase peptides are predicted to be immunogenic, and out of all 19 peptides, the top three predicted immunogenic peptides are all ZIKV NS3 helicase peptides (VLPEIVREA, AEMEEALRGL and TEVEVPERAW). Performing peptide sequence matching, we compare our 19 ZIKV peptides to all theoretical peptides of the human proteome to assess whether they may mimic endogenous human peptides. We identify 16 ZIKV peptides to have some degree of homology with human peptides (Table 5). Importantly, every comparison has a mismatch of at least two, and there are no more than seven matched amino acids in continuous order. As eight is the minimum peptide length for binding to the HLA-I groove, none of the identified ZIKV peptides mimics endogenous human peptides. To conclude, ZIKV NS3 helicase is a rich source of predicted immunogenic peptides, and the identified viral peptides do not mimic predicted peptides of the human proteome.

**Figure 4:**
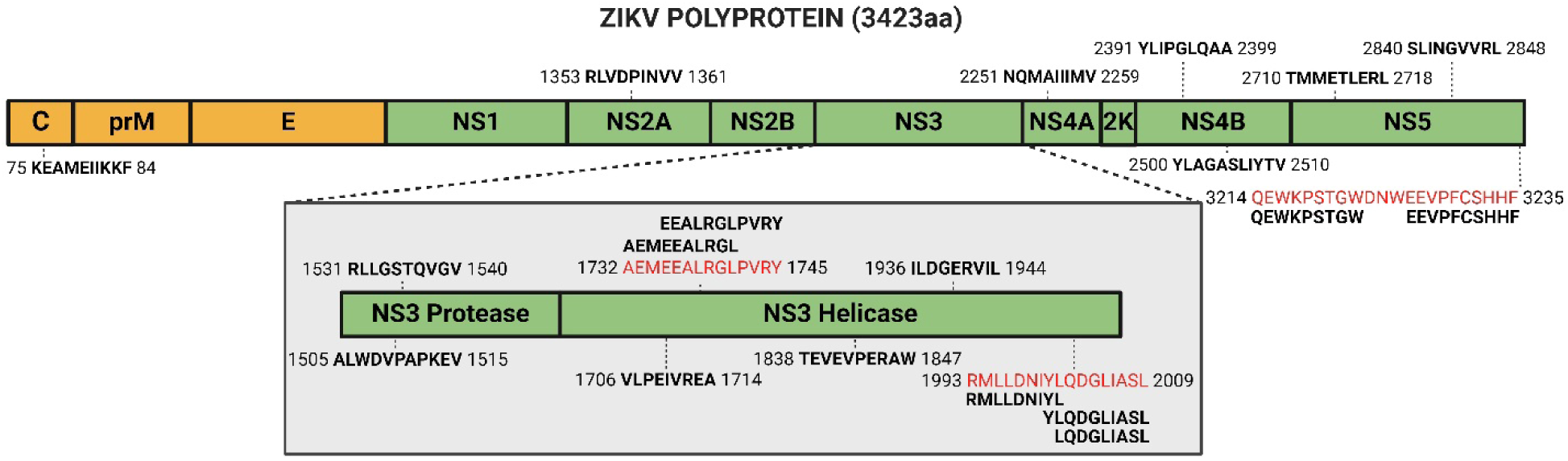
ZIKV polyprotein with mapped immunopeptides. The 19 identified ZIKV peptides are in bold, with the flanking numbers denoting the site in the polyprotein of the first and last amino acid. Peptide-rich sequences are indicated in red, with their corresponding peptides aligned above or below. ZIKV polyprotein not to scale. Figure created in Created in BioRender. BioRender.com/r34k740BioRender.com.

**Table 5:**
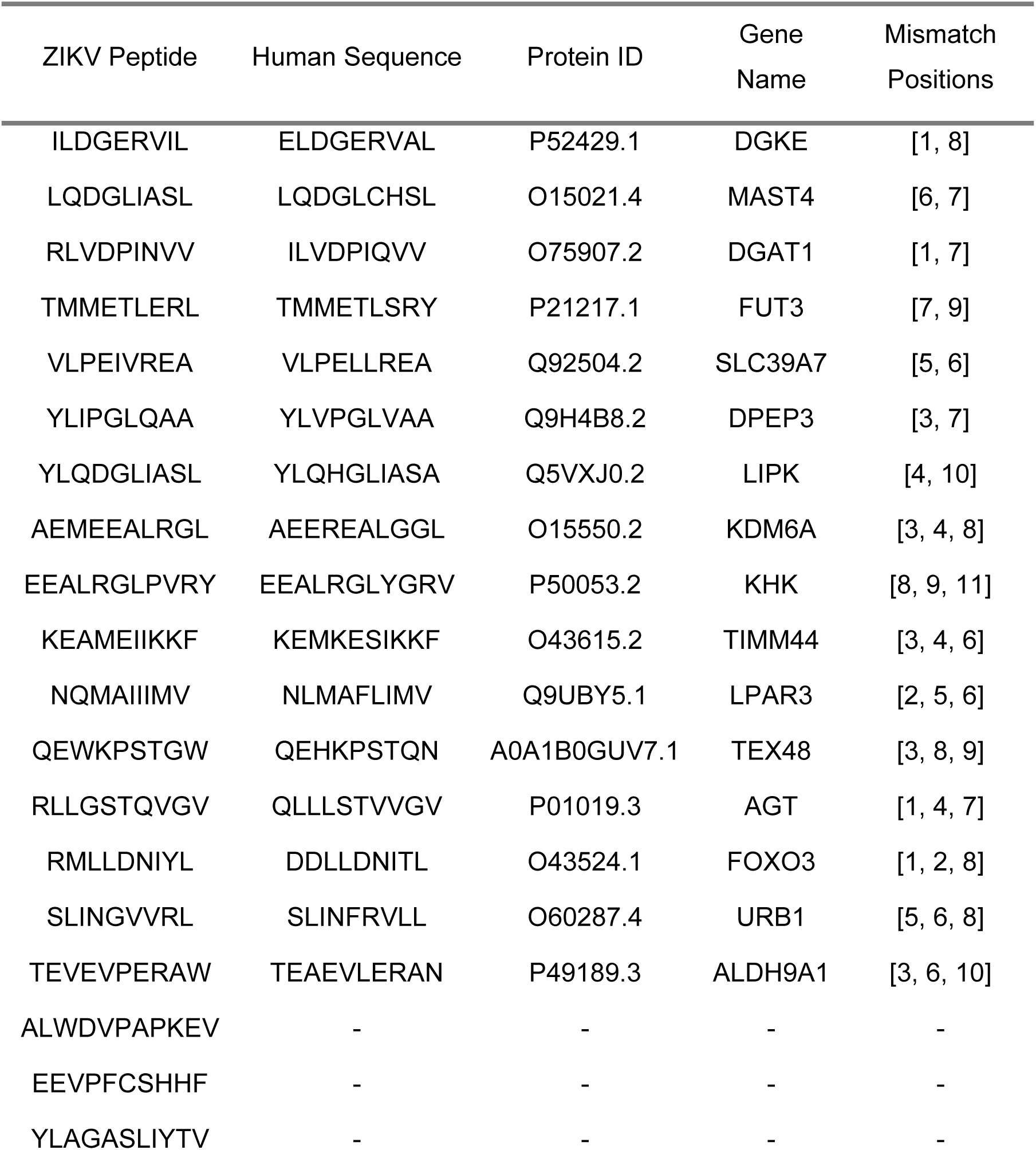
ZIKV : human peptide homology.

## Discussion

Downregulating HLA-I presentation is a method which cancer cells frequently utilise to help produce an immunosuppressive TIME (26). Here we investigate and show that ZIKV infection enriches the HLA-I pathway in both paediatric and adult brain tumour cells. Since brain tumour cells are non-professional antigen presenting cells, enrichment of HLA-I instead of HLA-II was expected. Supporting our observation of HLA-I pathway enrichment in brain tumour cells, HLA-A and HLA-B protein expression is upregulated in ZIKV-infected U251 GBM cells, where they act as a viral dependency factor and a regulator of cell viability in response to ZIKV infection, respectively (27). We propose that the enrichment of the HLA-I pathway following ZIKV infection of brain tumour cells may contribute to remodelling the TIME to make these commonly immunosuppressed tumours more immunogenic.

Here, we identify 19 HLA-I ZIKV peptides presented on the surface of USP7 brain tumour cells. Our work indicates the relevance of the HLA-I pathway and protein abundance to ZIKV peptide presentation and suggests a relationship as the most highly expressed viral proteins (NS3 and NS5) correlate with an increased number of presented peptides. The paediatric brain tumour cell line USP7 has highly advantageous traits for its use in our study because it possesses stem-like characteristics, it’s highly susceptible to ZIKV infection, and it’s homozygous for the globally most common HLA-A allotype HLA-A*02:01 (18). Of the 19 peptides observed here, twelve peptides are novel and seven have been previously observed elsewhere. Six of our HLA-A*02:01 peptides (RMLLDNIYL, YLQDGLIASL, ALWDVPAPKEV, YLIPGLQAA, SLINGVVRL and TMMETLERL) are presented on ZIKV-infected immortalised Priess B cells that are homozygous for HLA-A*02:01 (28). This supports our observations and indicates these six peptides as bona fide HLA-A*02:01 presented ZIKV epitopes. The peptides YLQDGLIASL, SLINGVVRL and AEMEEALRGL are recorded on the Immune Epitope Database (IEDB) under epitope IDs 2243385, 1311496 and 182464, respectively. NS3 YLQDGLIASL, NS4B YLIPGLQAA and NS5 SLINGVVRL stimulate memory T cell recall response in 57%, 14% and 57% of peripheral blood mononuclear cells (PBMCs) human patients previously infected with ZIKV (28,29). Additionally, SLINGVVRL was also one of the top dominant ZIKV epitopes in an immunocompetent HLA-A2 transgenic mouse model, capable of stimulating CD8+ T cells to produce IFNγ and TNFα (30). These observations validate our LC-MS/MS model as a viable approach to identify immunogenic HLA-I presented ZIKV epitopes. An inherent limitation of our work with non-professional antigen-presenting cells, is that we could not consider HLA-II-presented ZIKV peptides. There is great interest in performing this work as the infiltration and induction of CD4+ T cells contribute to ZIKV OV therapy efficacy (20,21). This would require feeding of infected cells to professional antigen presenting cells, such as dendritic cells, from which immunopeptidomes could then be captured as per (23).

It is important to understand if our ZIKV peptides have autoimmune implications by mimicking human peptides or if they may cross-react to stimulate memory T cells from previously encountered viral epitopes. Here, we show none of our ZIKV peptides to mimic human peptides, thus minimising the risk of autoreactive T cell activity and autoimmunity (31). THE HLA-I peptides ZIKV NS3 AEMEEALRGL and NS5 SLINGVVRL are homologous to a Dengue Virus (DENV) epitope and can stimulate memory cytotoxic T cells in Japanese Encephalitis Virus (JEV)-vaccinated HLA-A2 transgenic mice, respectively (30). Cytotoxic T cell depletion abolishes ZIKV OV efficacy, and long-term survivors of mouse glioma present with memory cytotoxic T cells that protect against tumour rechallenge (21). It is currently unknown if and how host humoral and cellular immunity to previous viral infection or immunisation may enhance or hinder ZIKV OV efficacy. We propose that brain tumour cell HLA-I presentation of a ZIKV epitope may enhance OV efficacy by co-opting a patient’s memory cytotoxic T cell immunity from a previous viral encounter against the tumour, and this deeply warrants investigation.

There are a multitude of complex interactions between an infected tumour and the immune system. Whilst both CD8+ and CD4+ T cells assist in GBM tumour clearance, myeloid cells protect GBM tumour cells from ZIKV infection through the secretion of type 1 interferons (32). Natural Killer (NK) cells can present with dichotomous function in response to OVs due to their contrasting antitumor and antiviral functions (33). Thus, it cannot be assumed that all ZIKV-stimulated immune cells will promote the therapeutic properties of ZIKV, and we must elucidate the roles and interplay between these cell types. Knowledge of the mechanisms of immune activation will factor into how ZIKV can be employed as an adjuvant for current immunotherapies. Thus far, ZIKV has proved an effective adjuvant to both immune checkpoint blockade (PD-1 and PD-L1) and vaccine-based immunotherapy to combat glioma and improve survival in mouse models (20–22). Our identification of novel HLA-I presented ZIKV epitopes contributes to the growing knowledge of how ZIKV can be employed as an immunotherapy and may assist in the development of novel epitope-specific immunotherapies against brain tumours.

Whilst our primary focus is to investigate the immunotherapeutic potential of ZIKV OV therapy, our research here is wider reaching, and has potential implications for (1) understanding fetal NPC depletion following ZIKV infection and (2) ZIKV epitope vaccination. CZS is primarily brought about by the depletion of fetal NPCs. ZIKV infection upregulates HLA-I processing and presentation in human NPCs at the transcriptome level and in microcephalic mouse brains at the transcriptome and proteome level (34,35). HLA-I presenting cells co-stain with infiltrating murine immune cells, resulting in neuronal cell death and microcephaly (35). Thus, cytotoxic CD8+ T cell clearance likely contributes to NPC depletion. To date, the immunopeptidome of ZIKV-infected NPCs is unknown, most likely due to the technical challenge of culturing NPCs to the high quantity required for immunopeptidomics. USP7 cells are of embryonal origin, closely resemble NPCs at the global gene expression level, and are immortalised so can be cultured to such quantities (18). As such, USP7 cells are an potential model to investigate the immunopeptidome of ZIKV-infected fetal NPCs. Interestingly, HLA-C is one of only 25 genes significantly upregulated in NPCs from CZS-affected twins compared to their unaffected dizygotic twin, thus possibly indicating HLA-I involvement in CZS development following congenital ZIKV infection (36). In the context of the developing fetus, we propose that the HLA-I peptides that we identify here may contribute to cytotoxic CD8+ T cell-mediated depletion of fetal NPC and subsequent CZS.

ZIKV exists as a single serotype and all strains could prove susceptible to a single immunogenic vaccine (37). Despite this, a vaccine is yet to be approved so ZIKV still poses a pregnancy risk and potential for re-emergence. Here, we identify ZIKV NS3 as a rich source of immunopeptides, producing over half of our observed epitopes. ZIKV NS3 peptides can promote NK cells and PBMC immune cell activity (28,39,40). ZIKV NS3 is the main antigenic T cell target, plays important roles during anti-ZIKV immunity, and a ZIKV NS3-based vaccine can stimulate the production of polyfunctional CD8+ T cells (38). Multiple *in silico* studies employ computational approaches to propose novel ZIKV epitope vaccines, one of which predicts the NS3 helicase sequence WLEARMLLDNIYLQDGLIASLYR as the richest ZIKV polyprotein epitope source (41–44). Our top identified region (RMLLDNIYLQDGLIASL) resides within this proposed vaccine. Additionally, we show all three ZIKV peptides within this region to be presented by the globally most common HLA-A*02:01 allotype, alongside identifying five more HLA-I peptides within the NS3 helicase domain. Thus, we confirm the ZIKV NS3 helicase as a rich epitopes source and promising vaccine candidate.

To summarise, immunopeptidomics is a potent and powerful tool to investigate viral peptide presentation to further our understanding of the immune responses orchestrated following ZIKV infection. Our results have possible future implications for the development of ZIKV OV therapy, epitope-specific immunotherapies or ZIKV epitope vaccine.

## Materials and Methods

### Cell Culture and ZIKV infection

USP7 cells were cultured under standard conditions (5% CO2, 37oC). Brazilian ZIKV KU365771 stocks were established by the Instituto Butantan (University of São Paulo, Brazil) in VERO cells and titrated by plaque-forming units (PFU) assay. For all infection experiments, USP7 cells were infected with ZIKV for 60 minutes prior to replacement with complete media. 24 hours post-infection (hpi) cells were collected, washed with PBS and stored as pellets at −80C. The infection experiments were performed in triplicate for the global proteome and once for the immunopeptidome. All controls were non-infected mock samples.

### Transcriptomics

#### HLA typing

High-quality total RNA was purified using the Monarch Total RNA Miniprep Kit (NEB #T2010S) as per the kit protocol from wildtype USP7 cells and sent to Novogene (UK) Company Limited for mRNA sequencing using the Illumina NovaSeq 6000 system (≥20 million 150 bp paired-end reads per sample). HLA typing was then performed using HISAT2 v2.2.0 with HISAT-genotype v1.3.0 in the default settings (45).

#### Adult glioma RNA-Sequencing Data Mining

RNA-Seq raw abundances of ZIKV-mCherry positive and ZIKV-mCherry negative primary cells isolated from eight GBM patients (IDs 42, 43, 45, 46, 50, 51, 54, 57) at 72hpi were downloaded from GEO (GSE178621) (32). The data was normalised using DESeq2, and GSEA was performed comparing the ZIKV-mCherry positive versus negative samples, as per the ZIKV-infected USP7 global proteome analysis (46,47).

### Proteomics

#### Global proteome sample preparation

50μg protein from mock and 24-hour infected USP7 cells (MOI 2) was mixed with 600μL methanol and 150μL chloroform for protein extraction. 450μL water was added to the sample, briefly vortexed and centrifuged at 14,000g for 5 min at room temperature (RT). The upper aqueous was removed and replaced with 450μL of methanol, the sample was then briefly vortexed and centrifuged again to pellet the proteins. The protein pellet was air-dried briefly before resuspension in 100μL 6M urea/50 mM Tris-HCl pH 8.0. The protein was reduced by 5 mM dithiothreitol for 30 min @37°C and alkylated by 15 mM iodoacetamide for 30 min @ RT in the dark. Protein was digested with 2μg trypsin/LysC mix (Promega) for 4h @37°C. 750μL 50mM Tris-HCl pH 8.0 was added, and the sample was incubated overnight @37°C. The digestion was terminated by the addition of 4μL TFA. The resultant peptide mixture was purified using HLB prime reverse phase μ-elution plates (Waters) by elution in 50μL 70% acetonitrile according to the manufacturers’ instructions and lyophilised. Peptides were reconstituted in 0.1% formic acid and applied to a Fusion LTQ orbitrap instrument set up as previously described.

#### LC-MS/MS analysis of global proteome

Tryptic peptides were reconstituted in 0.1% formic acid and applied to an Orbitrap Fusion Tribrid Mass Spectrometer with a nano-electrospray ion source as previously described. Peptides were eluted with a linear gradient of 3-8% buffer B (Acetonitrile and 0.1% Formic acid) at a flow rate of 300nL/min over 5 minutes and then from 8-30% over a further 192 minutes. Full scans were acquired in the Orbitrap analyser using the Top Speed data dependent mode, preforming a MS scan every 3 second cycle, followed by higher energy collision-induced dissociation (HCD) MS/MS scans. MS spectra were acquired at resolution of 120,000 at 300-1,500m/z, RF lens 60% and an automatic gain control (AGC) ion target value of 4.0e5 for a maximum of 100ms and an exclusion duration of 40s. MS/MS data were collected in the Ion trap using a fixed collision energy of 32% with a first mass of 110 and AGC ion target of 5.0e3 for a maximum of 100ms.

#### Data analysis for global proteome

Raw global proteome mass spec files were analysed using Peaks Studio 10.0 build 20190129 with spectra searched against the same database as used for immunopeptidomics. The false discovery rate (FDR) was estimated with decoy-fusion database searches and were filtered to 1% FDR. Relative protein quantification was performed using Peaks quantification module and normalized between samples using a histone ruler (48). Downstream analysis and visualizations were mainly performed in R using associated packages (49–52). GSEA analysis was performed on Log2(LFQ intensities) of mock and 24-hour infected USP7 cells (n=3) using a gene set database of Reactome innate and adaptive immune system pathways, nominal p-value of 0.05 adjusted for multiple hypotheses testing (FDR ≤ 0.25) (53). The top 50 and bottom 50 GSEA ranked proteins were plotted on a heatmap using GraphPad PRISM (10.0.3), with the ZIKV polyprotein being ranked and incorporated manually.

### Immunopeptidome analysis

#### Purification of HLA-I immunopeptides

Protein-A sepharose beads (Repligen, Waltham, Mass. USA) were covalently conjugated to 10 mg/mL W6/32 (pan-anti-HLA-I) or 5mg/mL HB145 (pan-anti-HLA-II) monoclonal antibodies (SAL Scientific, Hampshire, UK) using DMP as previously described (54). Frozen pellets of 1×10^8^ mock and ZIKV-infected USP7 cells (MOI 1) were re-suspended in 5mL of lysis buffer (0.02M Tris, 0.5% (w/v) IGEPAL, 0.25% (w/v) sodium deoxycholate, 0.15mM NaCl, 1mM EDTA, 0.2mM iodoacetamide supplemented with EDTA-free protease inhibitor mix), and rotated on ice for 30 min to solubilise. Homogenates were clarified for 10 min @2,000g, 4°C and then for a further 60 min @13,500g, 4°C. 2mg of anti-HLA-I conjugated beads were added to the clarified supernatants and incubated with constant agitation for 2h at 4°C. The captured HLA-I/*β*_2_microglobulin/immunopeptide complex on the beads was washed sequentially with 10 column volumes of low (isotonic, 0.15M NaCl) and high (hypertonic, 0.4M NaCl) TBS washes prior to elution in 10% acetic acid and dried under vacuum. Column eluates were diluted with 0.5 volumes of 0.1% TFA and then applied to HLB-prime reverse phase columns (Waters, 30mg sorbent/column). The columns were rinsed with 10 column volumes of 0.1% TFA and then the peptides were eluted with 12 sequential step-wise increases in acetonitrile from 2.5-30%. Alternate eluates were pooled and dried using a centrifugal evaporator and re-suspended in 0.1% formic acid.

#### LC-MS/MS analysis of HLA-I peptides

HLA peptides were separated by an Ultimate 3000 RSLC nano system (Thermo Scientific) using a PepMap C18 EASY-Spray LC column, 2μm particle size, 75μm x 50cm column (Thermo Scientific) in buffer A (0.1% Formic acid) and coupled on-line to an Orbitrap Fusion Tribrid Mass Spectrometer (Thermo Fisher Scientific, UK) with a nano-electrospray ion source. Peptides were eluted with a linear gradient of 3%-30% buffer B (Acetonitrile and 0.1% Formic acid) at a flow rate of 300nL/min over 110 minutes. Full scans were acquired in the Orbitrap analyser using the Top Speed data dependent mode, performing a MS scan every 3 second cycle, followed by higher energy collision-induced dissociation (HCD) MS/MS scans. MS spectra were acquired at resolution of 120,000 at 300 m/z, RF lens 60% and an automatic gain control (AGC) ion target value of 4.0e5 for a maximum of 100ms. MS/MS resolution was 30,000 at 100m/z. Higher-energy collisional dissociation (HCD) fragmentation was induced at an energy setting of 28 for peptides with a charge state of 2-4, while singly charged peptides were fragmented at an energy setting of 32 at lower priority. Fragments were analysed in the Orbitrap at 30,000 resolution. Fragmented m/z values were dynamically excluded for 30 seconds.

#### Data analysis for immunopeptidome

Raw spectrum files were analysed using Peaks Studio 10.0 build 20190129, with the data processed to generate reduced charge state and deisotoped precursor and associated product ion peak lists which were searched against a Uniprot database (20,350 entries, 2020-04) appended with the full sequences for ZIKV strain (Brazil KU365779.1; 2015): 10 entries. A contaminants list (245 entries) in unspecific digest mode was applied (55). Parent mass error tolerance was set a 5ppm and fragment mass error tolerance at 0.03 Da. Variable modifications were set for N-term Acetylation (42.01 Da), Methionine oxidation (15.99 Da) and carboxyamidomethylation (57.02 Da) of cysteine. A maximum of three variable modifications per peptide were set. The false discovery rate (FDR) was estimated with decoy-fusion database searches and were filtered to 1% FDR. The search results were further refined using the MS-Rescue package (56). Downstream analysis and visualizations were performed in R using associated packages (49–52). Peptide binding motifs were identified using unsupervised clustering methods MixMHCp2.1 and MoDec, for class I and class II HLA peptides, respectively (57,58). Peptide binding affinities predicted using NetMHC 4.0 and NetMHCIIpan 4.0 for class I and class II HLA peptides respectively (59–61). The IEDB T cell class I Immunogenicity (1.0) tool predicted ZIKV peptide: HLA-I complex immunogenicity, with selected settings of Peptide Length(s) 9-11-mer, MHC Allele(s) HLA-A*02:01 and HLA-B*44:02, and Allele Specific anchor positions (62). The IEDB PEPMatch (0.9) tool performed ZIKV peptide sequence matching against the human proteome, with the result specifying the best match per peptide and a maximum mismatch of three (63).

The mass spectrometry proteomics data have been deposited to the ProteomeXchange Consortium via the PRIDE partner repository with the dataset identifier PXD037627 and 10.6019/PXD037627 (64).

## Acknowledgments

RME acknowledges financial support for this project from UKRI (MR/S01411X/1), Wessex Medical Research Trust, Rosetrees Trust and The Little Princess Trust.

